# HPV16 Utilizes Phospholipase C(s) for its Genome Egress

**DOI:** 10.64898/2026.09.03.749318

**Authors:** Timothy R Keiffer, Abida Siddiqa, Mireya Represa-Perez, Hiba Zabir, Anand Kushwaha, Martin Sapp, Katarzyna Zwolinska

## Abstract

Nuclear delivery of human papillomavirus (HPV) requires infected cells to undergo mitosis. Incoming HPV DNA is protected by a transport vesicle structure, which becomes unstable post-mitosis. We have previously shown that HPV genome egress takes on average 4 hours post-mitosis completion; thus, we propose that an enzymatic process is involved in degrading this transport vesicle to allow genome egress. Several phospholipase C (PLC) isoforms, including PLCδ3, PLCζ1, and PLCL1, have been previously implicated in HPV infection in large-scale siRNA screens, and the parvovirus VP1 capsid protein has phospholipase enzymatic activity. Therefore, we hypothesized that cellular phospholipases residing in the nucleus mediate egress of the HPV genome from transport vesicles. We utilized siRNA-mediated knockdown and CRISPR/Cas9 knockout techniques to target specific PLC isoforms in both HeLa and HaCaT cell lines. We found that targeting PLC isoforms PLCβ1, β4, δ3, δ4, and γ1 significantly decreased HPV infection in both HeLa and HaCaT cells, as measured using a luciferase-based reporter assay. We also discovered a novel interaction between HPV16 minor protein L2 and PLCs β1, β4, δ1, δ3, and δ4. Furthermore, we observed that knockout of PLCβ4 and δ4 delayed egress of the HPV genome from nuclear membrane-bound vesicles after infection of HeLa cells. We propose that HPV utilizes phospholipase Cs to achieve genome egress from its protective vesicular structure after nuclear delivery.

**Importance:** The cellular factors involved in the later stages of HPV entry remain to be elucidated. Previous data gleaned from large-scale siRNA screens implicated the involvement of phospholipases in HPV infection. We expanded upon these findings and discovered that the individual phospholipase C (PLC) isoforms β4 and δ4 are important for HPV infectivity. We further linked the loss of infectivity in PLCβ4- and PLCδ4-deficient cells to a defect in HPV genome release from its transport vesicle, which we demonstrated using our differential staining technique. Furthermore, we discovered a novel interaction between the minor protein L2 of HPV16 and PLC isoforms β1, β4, δ1, δ3, and δ4.

## Introduction

Over 400 types of human papillomavirus (HPV) have been identified to date (1), and while most infections by these HPVs yield either asymptomatic infection or “common warts” (2–4), a subset of approximately 20 HPV subtypes has been characterized as “high-risk strains” due to their capability of causing anogenital, oropharyngeal, and cervical carcinomas (5–7). HPV subtypes 16 and 18 are high-risk strains, and these are responsible for over 70% of worldwide cervical cancers (8), along with the majority of other HPV-related cancers (9). HPV-associated oropharyngeal cancers are on the rise (10, 11). While prophylactic vaccines are available against several high-risk strains (12, 13), these will not address immediate- to intermediate-term HPV-related cancer burdens due to insufficient vaccination rates (14), suboptimal cervical cancer screening rates (15), and the latency between HPV infection and tumor detection (16). This warrants further investigation into the life cycle of HPV to discover novel druggable targets and stratagems. Our manuscript focuses on high-risk HPV subtype 16 (HPV16).

HPV16 virions are composed of L1 and L2, the major and minor capsid proteins respectively (17–19), that encapsulate the 8kb double-stranded DNA genome (20). HPV16 infects basal keratinocytes at mucosal membranes after first binding to the extracellular matrix (ECM) of the basement membrane (21–24). We have previously published several reviews focusing on this initial entry step of HPV16 (25, 26). Briefly, upon virion binding to ECM and during its subsequent endocytosis, a cascade of conformational changes occurs in the major and minor capsid proteins (27–32). These conformational changes allow the L2 protein to extend through the endosomal membrane upon acidification of HPV16 virion-containing endosomes (33–35), thus allowing L2 to be a substrate for various cytosolic proteins such as the retromer complex (36–41) for further trafficking of the HPV16 genome (42, 43). Mitosis is absolutely required as part of the viral lifecycle to allow the HPV viral genome (44, 45), protected from host sensor detection by its membrane-derived transport vesicle, to reach the host mitotic chromosomes (46–48).

While the exact composition of the HPV genome transport vesicle remains unknown, we can make a reasonable conjecture as to its probable makeup. Since HPV is trafficked to the trans-Golgi network (TGN) and then is directed towards the nucleus during the onset of mitosis (49–53), then at least a portion of the protective vesicle surrounding the HPV genome should be comprised of membranes derived from the Golgi apparatus. Vesicles derived from the Golgi can contain specific lipids such as phosphatidylinositol 4-phosphate (PI(4)P) (54) or specific proteins such as COPI (55), which allow the Golgi to act as a central sorting station for macromolecules in transit (56). Still, the exact nature of this HPV-derived vesicle remains a gap in our understanding of HPV initial entry.

There remain significant other gaps in the understanding of the latter part of initial HPV16 entry, especially regarding the cellular factors required for plus-end movement of the genome towards the nucleus from the microtubule-organizing complex (57), and how the HPV genome is ultimately released from its vesicle. Our current study aimed to elucidate how the HPV genome egresses from its protective vesicle post-nuclear delivery. We have previously found that HPV16 pseudogenome release from transport vesicles was delayed by an average of 4h compared with pseudogenome introduction via transfection (48), implying an enzymatic mechanism of HPV16 genome release. Also, transmission electron microscopy (TEM) studies have implied that HPV16 particles and their protective vesicles are unstable after post-mitotic nuclear membrane reformation (46). Given that several phospholipase C (PLC) isoforms, including δ3, ζ1, and L1, were previously implicated in HPV16 infection via large-scale siRNA screens (38, 44) and the aforementioned delay in genome accessibility, we hypothesized that host phospholipases, in particular, phospholipase C (PLC)s, are involved in HPV genome egress post-nuclear genome delivery. There is precedent for this, as the VP1 capsid protein of parvoviruses has phospholipase A2 activity, which is required for the incoming virus to escape the endosome (58). PLCs are capable of hydrolyzing the phosphodiester bond of phospholipids such as (mainly) phosphatidylinositol 4,5-bisphosphate (PI(4,5)P_2_), phosphatidylinositol 4-phosphate (PI(4)P), and phosphatidylinositol (PI); PLC cleavage of (PI(4,5)P_2_) yields diacylglycerol (DAG) and inositol 1,4,5-trisphosphate (IP₃) (59, 60). Several PLC isoforms, including β1, β4, δ1, δ4, and γ1 localize to the nucleus, although the δ1 isoform is known to shuttle between the cytoplasm and the nucleus (61–69).

Herein, we utilized luciferase construct-containing reporter HPV16 pseudoviruses to determine the effects of PLC disruption on HPV cell infectivity and HPV genome accessibility post-mitosis. We provide evidence via infectivity assays and confocal microscopy on cells subjected to our previously established differential staining protocol (48) that selective PLC knockout of isoforms β4 and δ4 decreases HPV infectivity and HPV genome accessibility. Finally, we propose several mechanisms by which PLCs might be recruited to HPV-containing vesicles post-mitosis.

## Results

### Select PLC Isoforms Required for Efficient HPV16 Infection

Given the availability of inhibitors against phospholipase A2 (ACA and PACOCF3, (70)) and phospholipase C (U73122) (71), we first wanted to determine if chemical inhibition of these phospholipases impeded infection in established HaCaT and HeLa cells. We detected a decrease, although not significant, of HPV16 infectivity in HaCaT cells treated with ACA 24hpi and 48hpi (Fig. 1A); no significant effect on HPV infectivity at either time point was noted in ACA-treated HeLa cells (Fig. 1B). PACOCF3 treatment significantly decreased HPV16 infectivity in both HaCaT (Fig. 1A) and HeLa (Fig. 1B) cells at the 48hpi time point. The phospholipase C inhibitor treatment (U73122) decreased HPV16 infectivity at both 24 and 48hpi time points in HaCaT cells (Fig. 1A), but not significantly at either time point in HeLa cells (Fig. 1B). Since we did not see an earlier HPV16 infectivity defect with the phospholipase A2 (PLA2) inhibitors, and inhibition of PLA can arrest cell division (72, 73), we decided to follow up more closely with the effects PLCs have on HPV16 infectivity.

**Figure 1.**
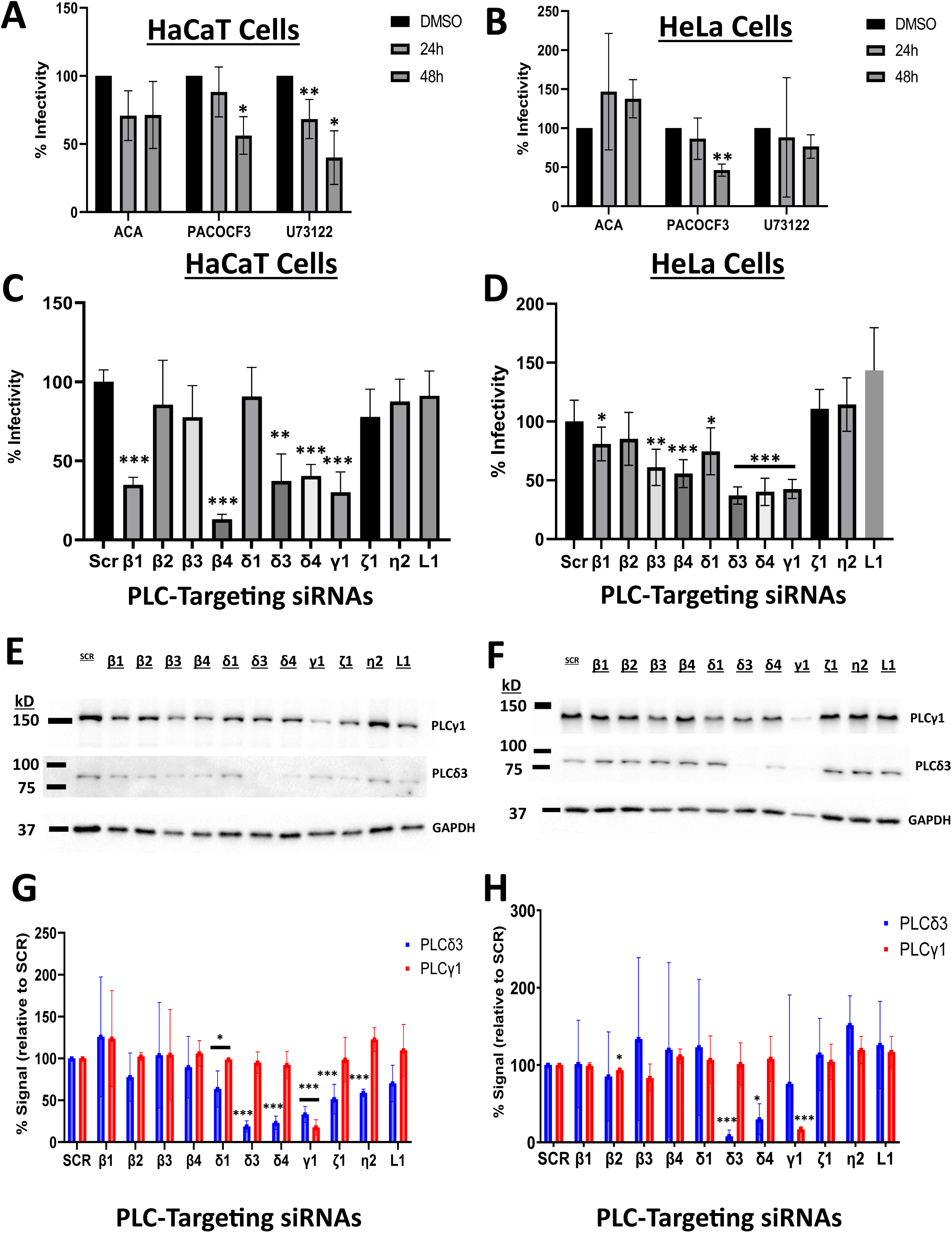
Chemical inhibition of and siRNA-mediated knockdown of phospholipase Cs (PLCs) decreased HPV16 infectivity of HeLa and HaCaT cells. (A-B) HaCaT (A) and HeLa (B) cells were infected with HPV16 pseudovirions at a viral genome equivalent (VGE) of approximately 100 in the presence of one of these compounds: N-(p-amylcinnamoyl) anthranilic acid (ACA), palmityltrifluoromet hylketone (PACOCF3), 1-[6-[[(17β)-3-Methoxyestra-1,3,5(10)-trien-17-yl]amino]hexyl]-1H-pyrrole-2,5-dione (U73122), or DMSO vehicle control. Infectivity of cell lines was determined by luciferase reporter assay 24h or 48h post-infection. Infectivity of cells treated with DMSO was set to “100% infectivity”, with infectivity of cells treated with other chemicals charted as percentages relative to DMSO control. Differences of infectivity values for each inhibition condition were analyzed using Student’s t-test (*N* = 3; *, P < 0.05; **, P < 0.01). (C-D) HaCaT (C) and HeLa (D) cells were first reverse-transfected with 20pmol of siRNAs either scrambled (SCR) control, or siRNAs targeting an individual PLC isoform (β1, β2, β3, β4, δ1, δ3, δ4, ζ1, η2, or L1); for the PLCγ1 (γ1) isoform, 10pmol of SCR or PLCγ1-targeting siRNAs were reverse-transfected into HaCaT (C) and HeLa (D) cells as with the other PLC-targeting siRNAs. Cells were then infected with HPV16 pseudovirions (VGE = 100) for either 48h (C, HaCaT cells) or 24 (D, HeLa cells). Infectivity of treated cells was then determined by luciferase reporter assay. Infectivity value differences for each siRNA treatment condition were analyzed using Student’s t-test (*N* = 3; *, P < 0.05; **, P < 0.01; ***, P < 0.001). (E-F) Western blots of HaCaT (E) and HeLa (F) lysates from cells treated with siRNAs used in (C & D); Westerns were probed with antibodies against PLCγ1, PLCδ3, and GAPDH. Blots are representative of 3 independent experiments. *N*=3. (G-H) Plots of PLCγ1 and PLCδ3 expression of Western blots (E-F), from either HaCaT (E) or HeLa (F) cells. Expression levels were normalized to GAPDH signal and charted relative to PLCγ1 and PLCδ3 signals in SCR siRNA-treated cells, which were plotted as “100% expression”. (*N*=3; *, P < 0.05; **, P < 0.01).

We next focused on determining the effect of targeting individual PLC isoforms on HPV infectivity, given the decrease in HPV16 infectivity observed in the HaCaT cells treated with U73122. U73122 has different inhibitory effects on different PLC isoforms, as U73122 does not inhibit PLCβ1, PLCβ3, or PLCβ4 isoforms (74). Also, there is no listed enzymatic activity data of U73122 on the PLCδ3 isoform itself. Therefore, we used siRNA-mediated knockdown of select PLC isoforms to determine the effect of their perturbation on HPV infectivity. Knockdown of PLC isoforms β1, β4, δ3, δ4, and γ1 significantly decreased HPV16 infectivity of both HaCaT cells (Fig. 1C) and HeLa cells (Fig. 1D). Additionally, PLCβ3 and δ1 knockdown decreased HPV infectivity in HeLa cells (Fig. 1D). We had difficulties using commercially available PLC antibodies to confirm siRNA-mediated PLC knockdown via Western blot, but we were able to narrow down the potential antibodies to use to demonstrate efficient siRNA knockdown to PLCβ4, PLCδ3, and PLCγ1-targeting antibodies. While using anti-PLCβ4 on these lysates gave us inconsistent nonspecific banding, we were able to show successful siRNA-mediated knockdown of the PLCδ3 and PLCγ1 isoforms by Western blot (Fig. 1E-1F). Given our PLC antibody troubles, we used PLCδ3 and PLCγ1 signal as proxies to determine the specificity of our PLC-mediated knockdowns. While we observed variable PLCδ3 decreases in HaCaT cells with knockdowns of PLC isoforms δ1, δ4, γ1, ζ1, and η2 (Fig. 1G), we only observed a concurrent PLCδ3 decrease in HeLa cells with knockdown of PLCδ4 (Fig. 1H). With regard to PLCγ1 signal, while we have statistical differences of this isoform in HaCaT and HeLa cells transfected with siRNA-targeting PLC isoforms δ1 and β2, respectively, versus control siRNA-treated conditions (Fig. 1G and 1H), the levels of PLCγ1 under these transfection conditions are still above 95%, thus there is no significant decrease of PLCγ1.

### Transfected HPV16-L2 Forms Complexes with PLCβ1, β4, δ1, δ3, and δ4

Due to the published findings that the minor protein of HPV16, L2, forms complexes with various host factors (75, 76), we decided to test if L2 could also potentially form complexes with PLCs. First, we transfected overexpression constructs for either PLCδ4 or γ1 along with L2 into HeLa:Cas9 cells. We co-immunoprecipitated (CoIP) PLCδ4 with L2, while we were unable to show clear PLCγ1 pulldown using L2 (Fig. 2). Since we were having issues with PLCβ4 overexpression in HeLa:Cas9 cells (data not shown), we switched our CoIP conditions to overexpress several PLCs, β1, β4, δ1, δ3, δ4, and L2 in 293TT cells and tested if we could CoIP L2 via PLCs. We successfully pulled down L2 using PLC isoforms β1, β4, δ1, δ3, and δ4 (Fig. 3).

**Figure 2.**
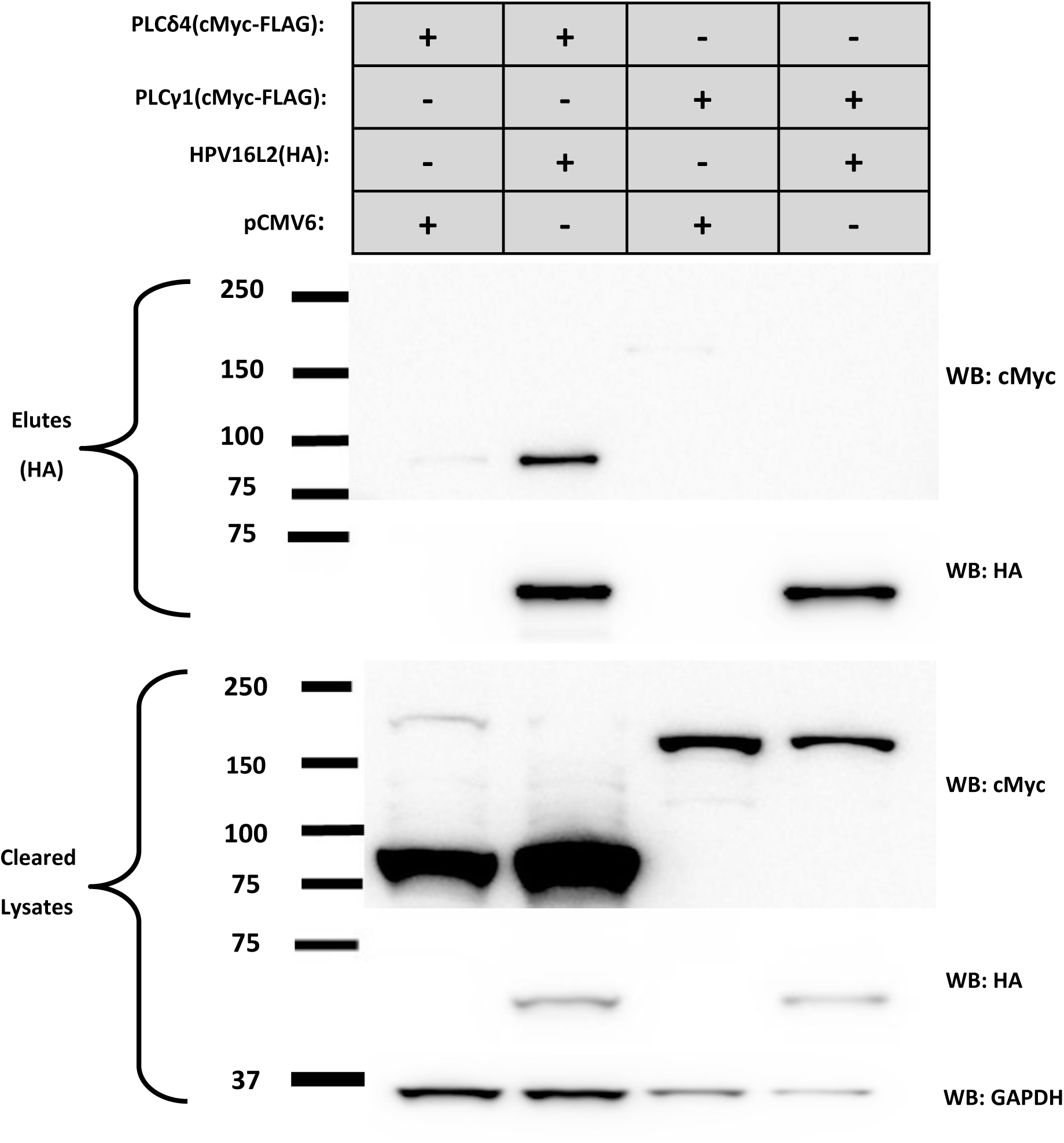
HPV16L2 forms complexes with phospholipase C (PLC) isoform β4 in HeLa:Cas9 cells. HeLa:Cas9 cells in 10cm plates were transfected with overexpression constructs for HPV16L2, encoding a C-terminal HA tag, and either an overexpression construct for PLCδ4 or PLCγ1, both containing the sequence for both C-terminal FLAG tag and c-Myc tag, or pCMV6, an empty vector control. Transfected cells were harvested 24h post-transfection, lysed in buffer containing 1% NP40 and 0.1% SDS, and then (cleared) lysates were incubated with magnetic beads coupled to anti-HA antibodies to pull down L2. Co-immunoprecipitation (CoIP) of PLCδ4 in bead elutions (“Elutes”) was determined using Western blot. Western blotting against the HA and c-Myc tags was used to confirm the presence of L2, PLCδ4, and PLCγ1 in “elutes” and “cleared lysates”. Blots are representative of three independent experiments (*N*=3).

**Figure 3:**
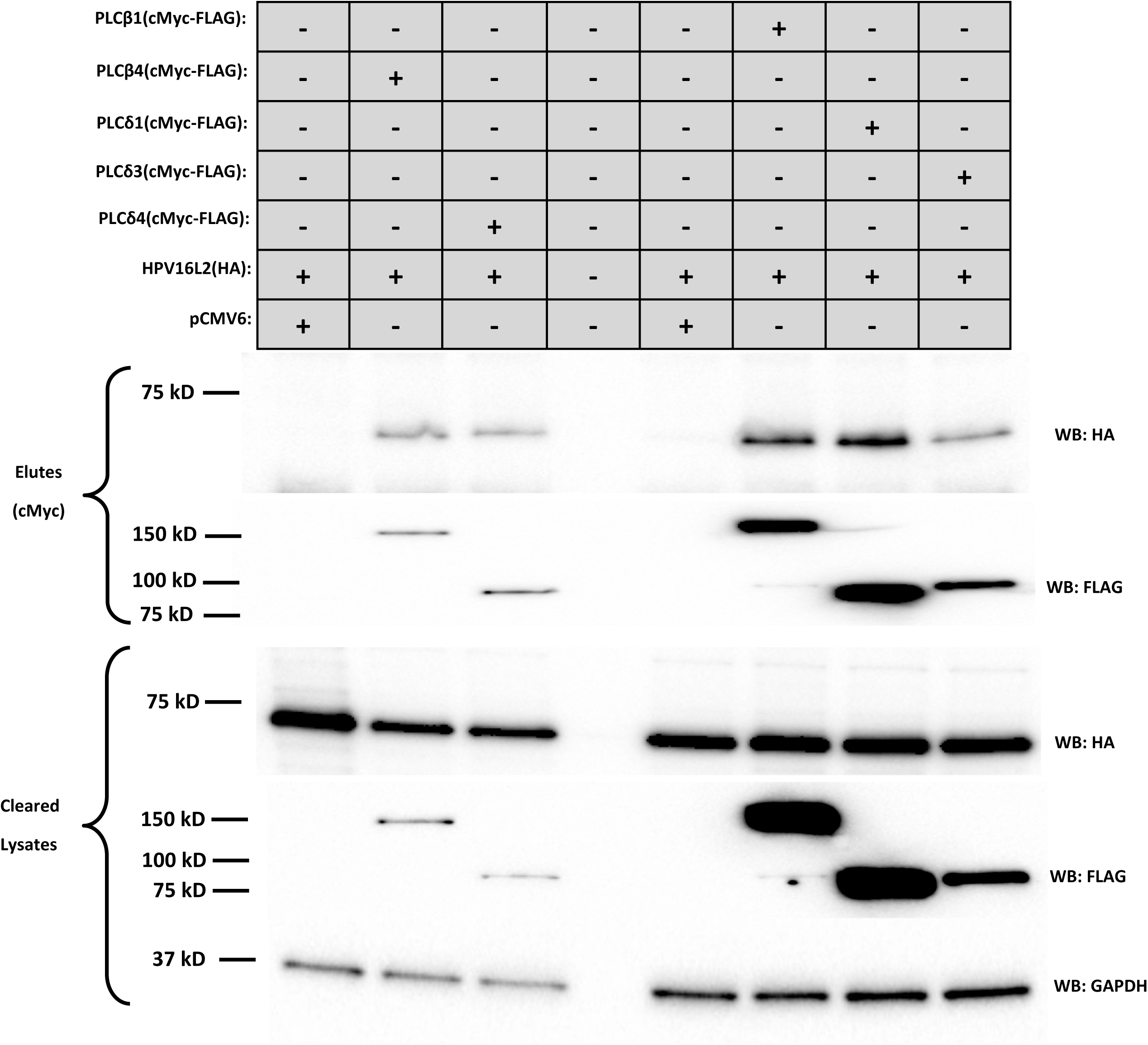
HPV16L2 forms complexes with phospholipase C (PLC) isoforms β1, β4, δ1, δ3, and δ4 in 2G3TT cells. 293TT cells in a 6-well format were transfected with overexpression constructs for HPV16L2, containing the C-terminal HA tag, and either an over-expression construct for PLC isoforms β1, β4, δ1, δ3, δ4, containing a C-terminal FLAG and a c-Myc tag, or the pCMV6 empty vector control. Transfected cells were harvested 24h post-transfection, lysed in buffer containing 1% NP40 and 0.1% SDS; cleared lysates were incubated with magnetic beads coupled to anti-c-Myc antibodies to pull down PLC isoforms. CoIP of L2 in “Elutes” was determined by Western blot; Western blotting against FLAG and HA tags was used to confirm the presence of select PLC isoforms and L2 in “elutes” and “cleared lysates”, respectively. Blots are representative of three independent experiments (*N*=3).

### Knockout of Select PLC Isoforms Decreases HPV16 Genome Egress from Transport Vesicles

While we wanted to utilize our differential staining technique to directly determine the effect of siRNA-mediated PLC knockdown on HPV16 genome accessibility, we encountered a major hurdle. Small amounts of cationic lipids, even from RNAiMax and similar reagents, can perturb the membranes that HPV utilizes during initial entry (77, 78). Due to this and our finding that knockdown of other PLC isoforms can affect PLCδ3 expression (Fig. 1G and 1H), we decided to utilize CRISPR/Cas9 to knock out the PLCβ4 or PLCδ4 genes in both HaCaT and HeLa cells and analyze genome accessibility in these cells. Due to a lack of suitable antibodies to confirm knockout of the PLCβ4 and PLCδ4 genes via Western blot, we used screening PCR to detect HaCaT (Fig. 4A) and HeLa (Fig. 4D) clonal cell lines containing edits to either the PLCβ4 or PLCδ4 gene. The PCR products from the unedited and edited HaCaT and HeLa cell lines were isolated from agarose gels, then sequenced and aligned to show the removal of DNA sections targeted by either the PLCβ4 or δ4 gRNAs. However, the expected PLCβ4 and PLCδ4 genome edits were clearly observed in the clonal HeLa:Cas9 cells but not in the HaCaT cells (Figs. 4E and 4B, respectively). The gRNA-mediated edits in the PLCβ4 gene of HeLa:Cas9 cells occurred in the C2-coding region, responsible for membrane-binding and catalytic core stabilization; the PLCδ4 gene edits occurred in the XY catalytic domain, which is conserved among the PLC family of enzymes and is indispensable for the main PLC enzymatic reaction of converting phosphatidylinositol 4,5-bisphosphate [PI(4,5)P_2_] into diacylglycerol (DAG) and inositol 1,4,5-trisphosphate (IP_3_) (79, 80).

**Figure 4.**
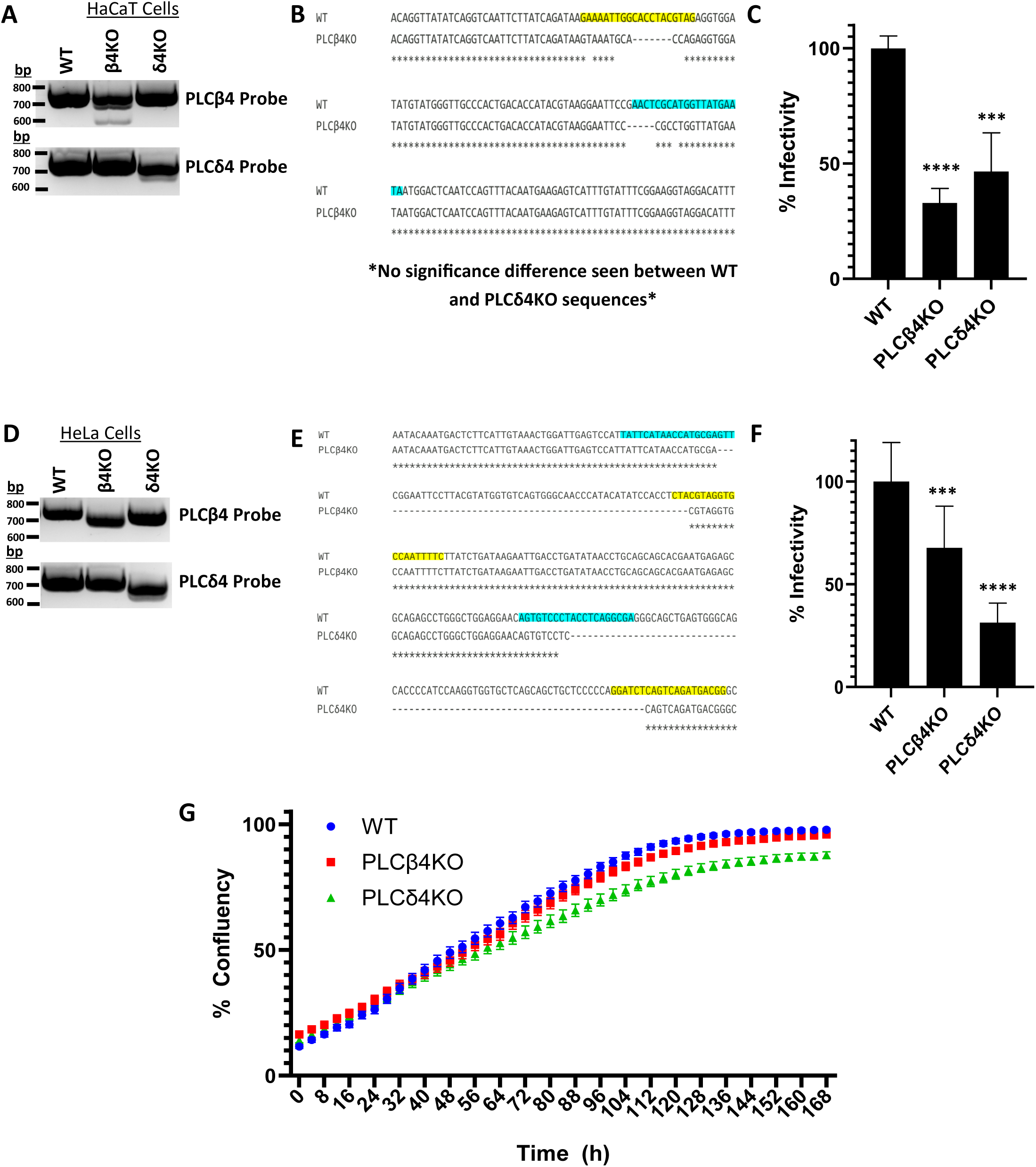
HPV16 infectivity is decreased in phospholipase C β4 (PLCβ4KO) and δ4 knockout (PLCδ4KO) cell lines. (A-D) HaCaT (A) and HeLa (B) cell lines were tested for edits to their PLCβ4 and PLCδ4 genes via PCR screening; total DNA was isolated from cell lines and probed via PCR to determine editing of either the PLCβ4 or PLCδ4 genes. To further confirm edits of either the PLCβ4 and PLCδ4 gene in select HaCaT (C) or HeLa (D) cells, PCR products were extracted from agarose gel and sequenced directly, with the sequences compared with wild-type PLCβ4 and PLCδ4 sequences. Sequences were aligned using Clustal Omega; sequences highlighted in yellow in the wild-type (WT) sequences were the gRNAs sequences used for the CRISPR/Cas9-mediated edits in the KO cell lines (E-F). Selected HaCaT (E) and HeLa (F) PLCβ4KO and PLCδ4KO cell lines were infected with 100 VGE of HPV16 pseudoviruses encapsulating a luciferase reporter; infectivities, as seen with luciferase activities, were analyzed 24h post-infection and compared with the wild-type (WT) parental (HeLa:Cas9) cells, which were set as “100% infectivity”. Infectivity value differences between WT, PLCβ4KO, and PLCδ4KO cells were analyzed using Student’s *t*-test (*N* = 3; ***, P < 0.001; ****, P < 0.0001). (G) The growth curves of the HeLa PLCβ4KO and PLCδ4KO cell lines, as compared to WT (HeLa:Cas9), were tabulated using IncuCyte (*N*=4).

Despite not detecting the full anticipated PLCβ4 or PLCδ4 gene edits in the clonal HaCaT cells (Fig. 4B), we tested the HPV16 pseudovirus infectivity of our selected clonal HaCaT and HeLa PLCβ4 or PLCδ4 knockout cells. All clonal cell lines tested had significantly less infectivity than the parental HeLa:Cas9 cells (Figs. 4C and 4F). To account for the possibility that the differences in infectivity were due to PLC gene disruption affecting cell proliferation, we analyzed cell growth between the HeLa:Cas9 parental and HeLa:Cas9-PLC-edited cell lines using Incucyte. No major difference was detected between the growth curves of parental or PLC-edited cell lines (Fig. 4G).

Next, we performed differential staining of infected parental and HeLa:Cas9 PLCβ4 and δ4KO cell lines – a brief summation of our technique is shown in Figure 5A. Both the PLCβ4 and δ4 knockout HeLa:Cas9 cell lines had significantly less EdU puncta visible after digitonin treatment in early interphase cells than the control cells (Fig. 5B). This corresponds to dramatically less HPV pseudogenome accessibility in both knockout cell lines (Fig. 5D). To ensure that the genome accessibility difference was not due to fewer HPV16 pseudogenomes entering the nucleus during mitosis, the total number of EdU puncta visible after Triton treatment in nuclei were counted. All cell lines had similar numbers of puncta (Fig. 5C).

**Figure 5.**
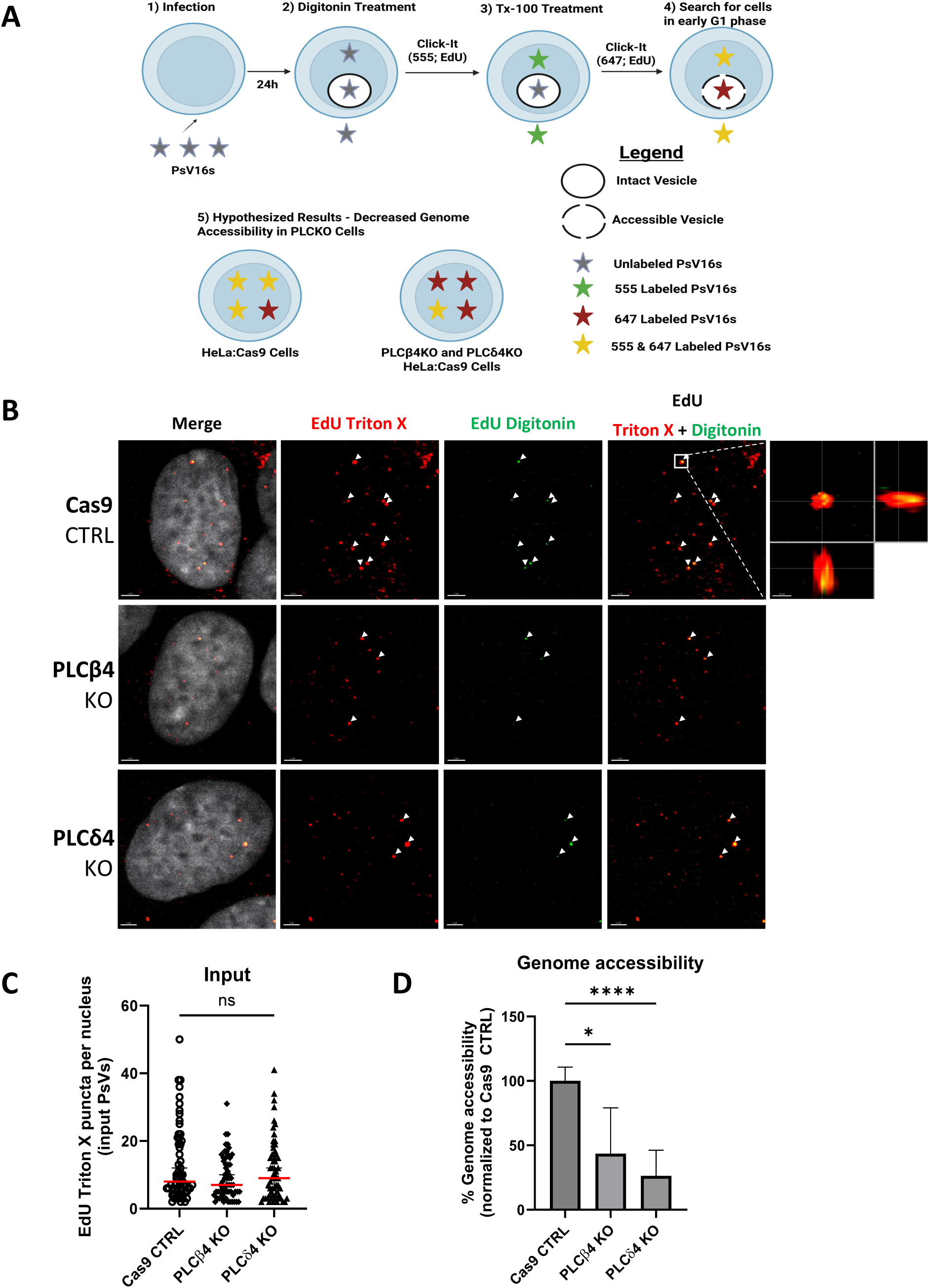
HPV16 genome accessibility is decreased in phospholipase C β4 knockout (PLCβ4KO) and δ4 knockout (PLCδ4KO) cell lines. A) Layout of the differential staining technique, based on a previously published protocol (113). Layout panel was generated using BioRender©. B) Representative images of HeLa:Cas9 (CTRL), PLCβ4KO, and PLCδ4KO cells 24hpi with 500 VGE of EdU-labeled HPV16 pseudoviruses after differential staining (56). 1st column (Merge) contains merged images of nuclear (Hoechst) staining with EdU staining. 2nd column (EdU Triton X) contains EdU staining images after Triton X-100 permeabilization (red); 3rd column (EdU Digitonin) contains EdU staining images after digitonin permeabilization (green). 4th column (EdU Digitonin + Triton X) consists of merged images of Triton X-100 (red) and digitonin permeabilization (green); white arrows show where both signals colocalize (yellow). A close-up of a cross-section in CTRL-infected cells showing a single locus with co-localized EdU signals after digitonin and TX-100 permeabilization (yellow) is shown in the far-right area of the row containing images from CTRL cells. This cross-section includes the transverse (big square), sagittal (small rectangle to the right), and frontal (rectangle below) view of the same 3D image. Image scale is 2µm. C) Quantification of nuclear HPV16 pseudovirus genomes per nucleus; EdU puncta after Triton-X permeabilization were quantified per nucleus. Median values with 95% CI, and data were analyzed by the Kruskal-Wallis test (*N* = 2; ns, not significant). D) Quantification of genome accessibility of infected CTRL, PLCβ4KO, and PLCδ4KO cells 24hpi; “genome accessibility” was quantified by adding the number of EdU Triton X-100 spots colocalized with EdU digitonin spots per cell and dividing that number by the number of total EdU Triton X-100 spots per cell. 100% genome accessibility was set to the median of Cas9 CTRL-infected HeLa Cells. Thirty early interphase cells per condition were quantified. Percent genome accessibility was plotted in GraphPad Prism and displayed as median values with 95% CI, with significance determined by Kruskal-Wallis test (*N* = 2; *, P < 0.05; ****, P < 0.0001).

## Discussion

This study aimed to address the question of how the HPV16 genome egresses from its protective transport vesicle that we have previously discovered indirectly via differential staining (48) and that other groups have visualized via TEM (46, 47). Given that previous large-scale siRNA screens implicated several phospholipases (PLCs) in HPV16 infection (38, 44), we expanded upon that by demonstrating that knockdown of several other PLC isoforms, including β1, β4, δ3, δ4, and γ1, significantly decreased HPV16 infectivity in both HaCaT and HeLa cells. We then decided to specifically target and analyze the role of PLC isoforms β4 and δ4 in HPV entry by knocking out these genes via CRISPR/Cas9 in HaCaT and HeLa cells. Clonal HaCaT and HeLa cell lines with disrupted PLCβ4 or PLCδ4 genes also had significantly decreased infectivity. HeLa:Cas9 cells with either disrupted PLCβ4 or PLCδ4 genes also had decreased HPV16 genome egress, lower than half of the amount seen in control cells.

This ability of incoming HPV16 to utilize cellular membranes to protect its genome via a unique transport vesicle, and then to usurp host enzymes to allow genome accessibility for transcription/translation, is yet another fascinating example of niche adaptation by viruses. Normally, dividing cells fragment their various compartments into vesicular structures to ensure equal distribution into daughter cells (81–83), but we are unaware of any normal condition where vesicles accumulate in the nucleus post-mitosis, presumably because such a defect would likely be deleterious to the cell due to interference with chromatin organization and increased nuclear envelope budding that is detected when the cell is stressed (84). However, herpesviruses utilize a unique vesicle-mediated nuclear egress mechanism to transport newly assembled capsids from the nucleus to the cytosol for their final maturation (85). Herpesviruses accomplish this via two conserved (among herpesviruses) proteins designated pUL31 and pUL34. These proteins form the core of the herpesvirus’ nuclear egress complex (86–90). Overexpression of pUL31 and pU34 from pseudorabiesvirus causes a buildup of vesicles in the perinuclear space in RK13 cells (91) and select mutations of pUL34 cause promiscuous budding of vesicles into the perinuclear space sans viral particles (92, 93).

It remains to be determined if HPV passively utilizes PLCs to aid in its genome egress or if HPV somehow directs select PLCs to its genome-containing vesicles. The latter is a distinct possibility due to the myriad interactions with host proteins mediated by L2 minor protein to facilitate HPV entry (36, 39, 43, 94, 95). Our discovery of this novel L2-PLC (β1, β4, δ1, δ3, δ4) interaction gives some credence to the possibility of L2-mediated direction of PLCs to HPV16-containing transport vesicles. However, we were unable to show that PLC(s) whose disruption did not affect genome egress (β1 and δ3, Fig 3B) also did not form complexes with L2, as both β1 and δ3 were able to pull down L2 in overexpression experiments (Fig 2B). This could be due to the transfected L2 not being processed in the same manner as L2 in HPV16 virions (27, 96–98) and thus L2 protruding from the HPV16 transport vesicle would have more limited accessibility to PLCs. It is also still unknown what binding motif(s) on PLCs HPV16 L2 would bind to, and vice versa. The middle T antigen protein of Merkel cell polyomavirus is phosphorylated by Src kinases on its SH2 motif, causing PLCγ1 recruitment to this protein and leading to an inflammatory signaling cascade (99). HPV16 L2 protein is phosphorylated at a conserved site (threonine 62) during infectious entry (100); however, this site is within the transmembrane-like domain (residues 45-67) and thus should be inaccessible for PLC binding (101). Oh *et al.* have recently shown through AlphaFold2 analysis and other mutational perturbations of HPV16 L2 that much of this protein is disordered (102) with no obvious motif for PLC binding. Another possibility is that L2 indirectly links PLCs towards HPV genome-containing vesicles via L2-PML(s) interaction. PML proteins allow HPV16 transcription initiation (103) and colocalize with HPV16 genomes prior to egress (104), demonstrating how HPV16 reorganizes these PML proteins for its own ends. PML nuclear bodies are incredibly dynamic with broad pleiotropy (105, 106), and published data show PML co-immunoprecipitation with nuclear PLCγ1, albeit in tested HCT116 cells but not in NB4 cells (107).

Additionally, the p62 protein has been shown to associate with PML proteins in HPV-infected cells (108), and there is one study showing that p62 and PLCγ1 can form complexes, albeit in mouse cells (109). We have observed potential co-staining of PLCβ4 and PLCγ1 with PML in HPV16 quasivirion-infected HaCaT cells (data not shown), despite our issues with antibody specificity, providing further support for this mechanism of bringing PLCs into proximity to HPV16 genome-containing vesicles for their dissolution. We summarize the possible ways PLCs can be brought into contact with the HPV genome-containing vesicle in Figure 6. Future research should focus on ascertaining the mechanism(s) by which PLCs are recruited to these HPV-containing vesicles, either by PLC interactions with PML, p62, the vesicle itself, or still-unknown host factors.

**Figure 6.**
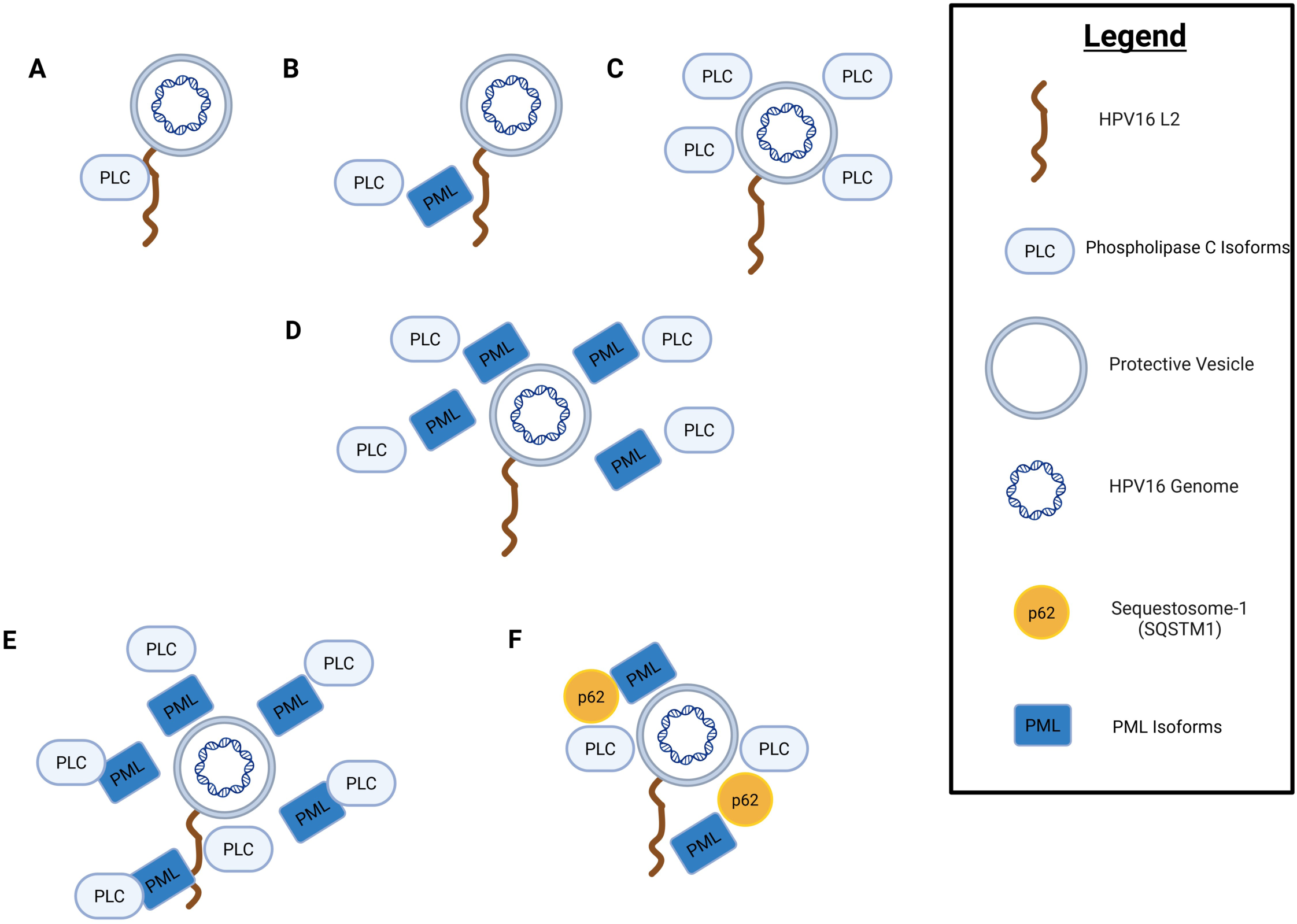
Proposed mechanisms of concentrating PLC activity against HPV genome-containing vesicles. A) PLCs nucleate around HPV16 L2 minor protein, directing PLC enzymatic activity toward the protective vesicle containing HPV16 genome. B and D) Select PML isoforms are brought in vicinity to the HPV genome-containing vesicle either through their interaction(s) with L2 protein or the vesicle itself, respectively. These incoming PMLs would then recruit PLC isoforms to themselves, thus concentrating PLC activity toward the protective vesicle. C) The accumulation of HPV16 genome-containing vesicles in HPV-infected cells could potentially be enough to nucleate nuclear-resident PLCs themselves on the vesicles. E) A Combination of PLC interactions with L2 and PML is required to release HPV genome from its vesicle. F) PML is recruited to either the vesicle or HPV16 L2, which then recruits p62, bringing PLCs and their enzymatic activity to the proximity of the HPV genome-containing vesicle. Figure was generated using BioRender©.

Given our data, we propose that we have discovered an uncharacterized “quality-control” mechanism by which vesicular structures that end up in the nucleus post-mitosis are dissolved by nuclear-migrating PLCs such as PLCβ4 and PLCδ4. Thus, it would behoove viruses such as HPV16 that require nuclear entry and mitosis as part of their lifecycle to take advantage of this proposed, little-known pathway. This needs to be elaborated further, as we still do not know all the factor(s) HPV16 needs for egress. Also, the details of the novel L2-PLC interaction, along with a potential PML protein linkage, require further clarification.

## Materials and Methods

### Materials

Phospholipase A (PLA) inhibitor compounds, anthranilic acid (N-(p-amylcinnamoyl); ACA), palmityltrifluoromethylketone (PACOCF3), and phospholipase C (PLC) inhibitor, 1-[6-[[(17β)-3-Methoxyestra-1,3,5(10)-trien-17-yl]amino]hexyl]-1H-pyrrole-2,5-dione (U73122), were all purchased from Enzo Life Sciences: BML-E178-0050, BML-ST336-0010, BML-ST391-0005, respectively. The following siRNA SMARTpools (ON-TARGETplus; a mixture of 4 siRNAs) used to target individual PLC isoforms for knockdown experiments were purchased from Horizon (Dharmacon); D-001810-10-05 (Non-targeting Pool), L-010280-00-0005 (Human PLCB1 (23236)), L-008274-00-0005 (Human PLCB2 (5330)), L-008485-00-0005 (Human PLCB3 (5331)), L-009594-00-0005 (Human PLCB4 (5332)), L-009149-02-0005 (Human PLCD1 (5333)), L-008649-01-0005 (Human PLCD3 (113026)), L-005065-01-0005 (Human PLCD4 (84812)), L-027529-01-0005 (Human PLCH2 (9651)), and L-009866-02-0005 (Human PLCL1 (5334)), L-008303-01-0005 (Human PLCZ1 (89869)). Additional siRNAs targeting PLCγ1 were purchased from Sigma-Aldrich, batch numbers WD12460035-WD12460036.

Plasmid constructs containing the guide RNAs (gRNAs) used to target PLC isoforms via CRISPR/Cas9 were purchased from VectorBuilder; pLV[2gRNA]-Neo-U6 was the construct backbone. The gRNA sequences used to target the PLCβ4 gene isoform were: GAAAATTGGCACCTACGTAG [gRNA#33313] & TATTCATAACCATGCGAGTT [gRNA#33323], and the gRNA sequences used to target the PLCδ4 gene isoform were: CCGTCATCTGACTGAGATCC [gRNA#3090] & AGTGTCCCTACCTCAGGCGA [gRNA#3107]. The primer sequences used to screen DNA isolated from individual cell lines for PLCβ4 gene edits were: 5’ - GGCCGTGGACACTATTTGTA – 3’ and 5’ – GGGTAGGGAAGAACTCTCTAGT -3’; primer sequences used to screen DNA for PLCδ4 gene edits were: 5’ – TGGCACAGGTATGAGATTTAGG -3’ and 5’ – GTTATGGGAGAGGTTGGGATTAT -3’.

Constructs encoding cDNA to overexpress C-terminally tagged FLAG PLCβ4 (Origene, RC217903) and PLCδ4 (Origene, RC202935), along with empty vector pCMV6 (Origene, SKUPS100001) were purchased commercially. The antibodies used for Western blots were anti-PLCδ3 (Origene, AP53337PU-N), anti-PLCγ1(Cell Signaling Technology, #2822S), anti-HA (Sigma-Aldrich, H9658), anti-DDK (OriGene, TA50011), anti-GAPDH (Cell Signaling Technology, #2118), and anti-mouse and anti-rabbit horseradish peroxidase-coupled secondary antibodies (Jackson ImmunoResearch Laboratories, 115-035-003 and 111-035-003, respectively).

#### Cell Culture Conditions

Standard HEK293TT and HeLa cells were cultured in DMEM (Corning, 10-013-CM) supplemented with 10%, L-Glutamax (ThermoFisher, 35050061), MEM non-essential ammo acids (ThermoFisher, 11140050), antibiotic/antimycotic (ThermoFisher, 15240062), and 2.5mg/ml Plasmocin® (InvivoGen, ant-mpp), while standard HaCaT cells were cultured in low-glucose DMEM (Corning, 10-013-CM) supplemented with 5% FBS (Atlanta Biologicals, S11150). HeLa:Cas9 cells were cultured in the same media as standard HeLa cells, with the additional supplement of 10µg/ml blasticidin (Invivogen, ant-bl-05). Clonal HaCaT cells containing PLC-targeting gRNAs were grown in same media as standard HaCaT cells, with the additional supplements of 10µg/ml puromycin (Sigma-Aldrich, P8833) and 400ng/ml G418 (Sigma-Aldrich, G8168), with clonal HeLa:Cas9 cells containing PLC-targeting gRNAs cultured in standard media supplemented with blasticidin and G418.

### siRNA-mediated knockdown of phospholipase C (PLC) isoforms

PLC isoform(s) expression was knocked down for infection experiments using 20pmol of scrambled (SCR) or PLC-targeting siRNA pools that were reverse-transfected into approximately 80,000 HeLa or HaCaT cells using Lipofectamine™ RNAiMAX Transfection Reagent (ThermoFisher, 13778030).

### Generation of HPV16 pseudoviruses and lentiviruses encoding PLC-targeting gRNAs

HPV16 pseudoviruses were generated as previously described (110–113). Briefly, 293TT cells were transfected with pShell16, encoding HPV16 L1 and L2, along with a reporter gene, either pGL3 (Promega), encoding a luciferase reporter, or pFWB, encoding green fluorescent protein (generously provided by Dr. John Schiller) using MATra A transfection reagent (IBA Lifesciences, 7-2001-100); HPV16 pseudovirions were then matured as previously described and isolated using OptiPrep Density Gradient Medium (Sigma-Aldrich, D1556) via ultracentrifugation for 3.5h at 50,000rpm at 16°C. HPV16 pseudovirions slated for use in pseudogenome-staining experiments had the thymidine analog EdU (5-Ethynyl-2′-deoxyuridine; Invitrogen, A10044) incorporated into their reporter genomes by supplementing the growth media of transfected 293TT cells with EdU (50µM) roughly 6h and 30h post-transfection. For the PLC isoform knockout, PLCβ4 and PLCδ4 isoform-targeting vectors were packaged into lentiviruses via transfection of 293TT cells using MATra-A reagent via psPAX2 and pMD2.G constructs as previously described (114).

### Generation of clonal PLC gene isoform knockout HaCaT and HeLa cell lines

We utilized the services of the LSU Health Shreveport Redox Molecular Signaling Core to generate HaCaT cells containing individual knockouts (KO) of the genes encoding either PLCβ4 or PLCδ4 isoforms; the Redox Core transduced Cas9 and PLCβ4- or PLCδ4-targeting gRNA-containing vectors into HaCaT cells. Given the availability of Cas9-expressing HeLa cells (HeLa:Cas9 cells), we transduced these cells with lentiviruses, generated using the psPAX2/pMD2G system, containing constructs encoding either PLCβ4- or PLCδ4-targeting gRNAs. Clonal HaCaT and HeLa cell lines were then isolated from the bulk population(s) by low-density seeding (100 cells/dish) into 150mm dishes, with the genomic DNA from individual clone(s) isolated using a NucleoSpin Blood, Mini Kit (Macherey-Nagel, 740951). PLCβ4 or PLCδ4 gene isoform edits from isolated clonal cell DNA were then screened via PCR using primers flanking the PLCβ4 or PLCδ4 gene sequences targeted by gRNAs. PLCβ4 and PLCδ4 gene edits in selected clonal HeLa cells were further confirmed via sequencing with Plasmidsaurus sequencing services using Oxford Nanopore Technology.

### Incucyte Analysis of Cell Growth

Parental HeLa:Cas9 and clonal HeLa:Cas9 cells with edited PLCβ4 and PLCδ4 genes were seeded on 4 wells of a 12-well plate, approximately 30,000 cells each, at a confluency of approximately 10%. The scan parameters of the Incucyte machine were set to capture images at 10x magnification of cell confluency every 4h up to 160h.

### Infectivity Assays

For infectivity assays with PLA and PLC chemical inhibitors, HaCaT or HeLa cells were first seeded into 12-well plates and then infected with HPV16 pseudoviruses encapsulating the luciferase reporter plasmid pGL3, at a viral genome equivalent (VGE) of 100. The vehicle control DMSO or inhibitor compound(s), 5µM each, were added at the same as pseudoviruses. Infected HeLa or HaCaT cells were then harvested either 24hpi or 48hpi, respectively, via trypsinization and transferred to Nucleon^TM^ (Thermo Fisher, 13610) 96-well plate(s). ONE-Glo™ Luciferase reagent (Promega, E6110) was used to lyse cells and the subsequent luciferase activity of lysates was quantified using a Tecan Spark plate reader and expressed as relative luciferase units (RLUs). “100%” infectivity was set to luciferase activity in control cells (DMSO-treated), with the “%” infectivity of other cell conditions plotted relative to control condition.

For infectivity assays with siRNA-treated HaCaT or HeLa cells or clonal cell lines, clonal cells were first seeded into 12-well plates 24h prior to infection or, for the siRNA-treated HaCaT or HeLa cells, they were first reverse-transfected with siRNAs also 24h prior to infection. Cells were then infected with HPV16 pseudoviruses at a VGE of 100, harvested, and had their luciferase activities ascertained the same way previously described for the inhibitor infectivity assays. “100%” infectivity for these infectivity assays was set to luciferase activity in control cells (such as cells treated with scrambled siRNAs, or parental cell lines), with the “%” infectivity of other cell conditions plotted relative to control conditions.

### Differential Staining

The full differential staining protocol has been previously described in DiGiuseppe *et al.* (115). Briefly, HeLa cells were infected with EdU-labeled HPV16 pseudoviruses containing pFWB reporter genome at a VGE of approximately 500 and then subjected to differential staining protocol at approximately 24hpi. Infected cells were partially permeabilized with digitonin (0.625µg/ml), with any accessible EdU-incorporated HPV pseudogenome labeled with AF555 dye using Click-iT reaction (Click-iT EdU imaging kit; Invitrogen, C10338); cells were then totally permeabilized with 0.5% TX-100, and all pseudogenomes were labeled with AF647 dye using a second Click-It reaction (Click-iT EdU imaging kit; Invitrogen, C10340). Confocal images from multiple fields per condition were obtained as Z-stacks with a Leica TCS SP5 confocal laser scanning microscope. For image analyses, early interphase cells were selected based on the number of nucleoli (>7). Quantification of genome accessibility was performed using Imaris 10.2.0 software, rendering EdU puncta as spots and quantifying colocalization of EdU puncta after permeabilization with Triton X-100 (TX-100) and digitonin (DIG). Genome accessibility was calculated as the number of EdU puncta after TX-100 that co-localized with EdU puncta after DIG permeabilization (accessible viral genomes), normalized to the total number of EdU puncta after TX-100 (total viral genomes) within the nucleus, and represented as a percentage. Imaging and analysis settings were maintained for each individual experimental replicate.

### Co-immunoprecipitations

We used both HeLa:Cas9 cells and 293TT cells for our co-immunoprecipitation (CoIP) experiments. For the HeLa:Cas9 cells, cells were transfected in 100cm dish format, at approximately 60% confluency, with 8.6 µg or 4.3µg, respectively, of plasmid constructs encoding either PLCδ4 or PLCγ1 (double-tagged with C-terminal FLAG and cMyc), and 8.6µg of either HA-tagged HPV16 L2 or pCMV6 (empty vector) constructs using 17.2µl of MATra-A transfection reagent per transfection. Transfected cells were harvested 24h post-transfection and lysed in 50mM Tris, 50mM NaCl, 2mM EDTA, pH 8.0 buffer containing 1% NP40 with 0.1% SDS, supplemented with protease inhibitors (cOmplete™, Mini, Roche, 11836153001). Cell lysates were cleared via centrifugation and then incubated with anti-HA magnetic beads (Pierce^TM^, 88836) end-over-end overnight at 4°C. Beads were then extensively washed with lysis buffer and proteins were eluted from them via boiling for 10min at >95°C. For 293TT transfections, cells were transfected in 6-well plate format, at approximately 60% confluency, with either 1.5µg of double-tagged (cMyc and FLAG) PLC-expressing constructs β1, β4, δ1, δ3, or δ4 and 1.5µg of HA-tagged HPV16-L2 or pCMV6 constructs. Transfections were again done with MATra-A transfection reagent, 3µl per transfection, and cells were harvested 24h post-transfection. Cells were lysed, and lysates were cleared the same way as with the HeLa cells, only this time the cleared lysates were incubated with anti-cMyc beads (MedChemExpress, HY-K0206) end-over-end for 1h at 4°C. Anti-cMyc beads were washed, and protein eluates were isolated from the beads in the same way as with the HeLa cells. The presence of PLC(s) and L2 in the cleared lysates and elutes was confirmed by Western blot; samples were separated on 4-20% gradient gels (Novex^TM^, Invitrogen, XPO4202) via SDS-PAGE, transferred to nitrocellulose membrane, and probed with aforementioned anti-FLAG (for PLC(s)) and anti-HA (for L2) antibodies, with chemiluminescent signal(s) visualized using West Pico substrates (ThermoScientific, 1863098 and 1863099).

## Acknowledgments

This research was funded by the National Institute of Health (NIH), grant numbers R01 A1164683, P20GM134974. PLC knockout HaCaT cells were generated using the services of the LSU Health Shreveport Redox Molecular Signaling Core (RRID:SCR_024778). HeLa cells expressing Cas9 (HeLa:Cas9) were graciously provided by Dr. Ana-Maria Dragoi of LSU Health Shreveport. The Incucyte machine was housed in the Innovative North Louisiana Experimental Therapeutics program (INLET) Core at LSU Health Shreveport (RRID:SCR_024990). Our confocal images were generated and analyzed using the instruments and services at The Microscopy Imaging Core at LSU Health Shreveport - Research Core Facility (RRID:SCR_024775). The Bioinformatics and Modeling Core (RRID:SCR_024779) supported the BioRender license. Mireya Represa-Perez is a fellow supported by the Doctoral Scholarship Program of the Barrie Foundation and LSU Health Shreveport. Lastly, we would like to acknowledge the late Dr. Martin J. Sapp, as this work would not have been initiated or even finished without his guidance and resources.

## References

1. McBride AA. 2022. Human papillomaviruses: diversity, infection and host interactions. Nat Rev Microbiol 20:95–108.

2. Al Aboud AM, Nigam PK. 2026. WartStatPearls. StatPearls Publishing, Treasure Island (FL).

3. Burd EM. 2003. Human papillomavirus and cervical cancer. Clin Microbiol Rev 16:1–17.

4. Workowski KA, Berman S, Centers for Disease Control and Prevention (CDC). 2010. Sexually transmitted diseases treatment guidelines, 2010. MMWR Recomm Rep Morb Mortal Wkly Rep Recomm Rep 59:1–110.

5. Baba SK, Alblooshi SSE, Yaqoob R, Behl S, Al Saleem M, Rakha EA, Malik F, Singh M, Macha MA, Akhtar MK, Houry WA, Bhat AA, Al Menhali A, Zheng Z-M, Mirza S. 2025. Human papilloma virus (HPV) mediated cancers: an insightful update. J Transl Med 23:483.

6. Cogliano V, Baan R, Straif K, Grosse Y, Secretan B, El Ghissassi F, WHO International Agency for Research on Cancer. 2005. Carcinogenicity of human papillomaviruses. Lancet Oncol 6:204.

7. Jain M, Yadav D, Jarouliya U, Chavda V, Yadav AK, Chaurasia B, Song M. 2023. Epidemiology, Molecular Pathogenesis, Immuno-Pathogenesis, Immune Escape Mechanisms and Vaccine Evaluation for HPV-Associated Carcinogenesis. Pathogens 12:1380.

8. Tommasino M. 2014. The human papillomavirus family and its role in carcinogenesis. Semin Cancer Biol 26:13–21.

9. Wei F, Georges D, Man I, Baussano I, Clifford GM. 2024. Causal attribution of human papillomavirus genotypes to invasive cervical cancer worldwide: a systematic analysis of the global literature. Lancet 404:435–444.

10. Aupérin A. 2020. Epidemiology of head and neck cancers: an update. Curr Opin Oncol 32:178–186.

11. Oral Cavity and Oropharyngeal Cancer Key Statistics. https://www.cancer.org/cancer/types/oral-cavity-and-oropharyngeal-cancer/key-statistics.html. Retrieved 5 April 2026.

12. Ghosh A, Chatterjee S, Dawn A, Das A. 2025. HPV Vaccines - An Overview. Indian J Dermatol 70:188–200.

13. Zhai L, Tumban E. 2016. Gardasil-9: A global survey of projected efficacy. Antiviral Res 130:101–109.

14. Jensen JE, Becker GL, Jackson JB, Rysavy MB. 2024. Human Papillomavirus and Associated Cancers: A Review. Viruses 16:680.

15. Liu X, Ning L, Fan W, Jia C, Ge L. 2024. Electronic Health Interventions and Cervical Cancer Screening: Systematic Review and Meta-Analysis. J Med Internet Res 26:e58066.

16. Doorbar J. 2023. The human Papillomavirus twilight zone – Latency, immune control and subclinical infection. Tumour Virus Res 16:200268.

17. Buck CB, Cheng N, Thompson CD, Lowy DR, Steven AC, Schiller JT, Trus BL. 2008. Arrangement of L2 within the papillomavirus capsid. J Virol 82:5190–5197.

18. Chen J, Wang D, Wang Z, Wu K, Wei S, Chi X, Qian C, Xu Y, Zhou L, Li Y, Zhang S, Li T, Kong Z, Wang Y, Zheng Q, Yu H, Zhao Q, Zhang J, Xia N, Li S, Gu Y. 2023. Critical Residues Involved in the Coassembly of L1 and L2 Capsid Proteins of Human Papillomavirus 16. J Virol 97:e0181922.

19. Chen XS, Garcea RL, Goldberg I, Casini G, Harrison SC. 2000. Structure of small virus-like particles assembled from the L1 protein of human papillomavirus 16. Mol Cell 5:557–567.

20. Mirabello L, Clarke MA, Nelson CW, Dean M, Wentzensen N, Yeager M, Cullen M, Boland JF, NCI HPV Workshop, Schiffman M, Burk RD. 2018. The Intersection of HPV Epidemiology, Genomics and Mechanistic Studies of HPV-Mediated Carcinogenesis. Viruses 10:80.

21. Bravo IG, de Sanjosé S, Gottschling M. 2010. The clinical importance of understanding the evolution of papillomaviruses. Trends Microbiol 18:432–438.

22. Culp TD, Budgeon LR, Christensen ND. 2006. Human papillomaviruses bind a basal extracellular matrix component secreted by keratinocytes which is distinct from a membrane-associated receptor. Virology 347:147–159.

23. Gheit T. 2019. Mucosal and Cutaneous Human Papillomavirus Infections and Cancer Biology. Front Oncol 9:355.

24. Selinka H-C, Florin L, Patel HD, Freitag K, Schmidtke M, Makarov VA, Sapp M. 2007. Inhibition of transfer to secondary receptors by heparan sulfate-binding drug or antibody induces noninfectious uptake of human papillomavirus. J Virol 81:10970–10980.

25. DiGiuseppe S, Bienkowska-Haba M, Guion LG, Sapp M. 2017. Cruising the cellular highways: How human papillomavirus travels from the surface to the nucleus. Virus Res 231:1–9.

26. Keiffer TR, Soorya S, Sapp MJ. 2021. Recent Advances in Our Understanding of the Infectious Entry Pathway of Human Papillomavirus Type 16. Microorganisms 9:2076.

27. Bienkowska-Haba M, Patel HD, Sapp M. 2009. Target cell cyclophilins facilitate human papillomavirus type 16 infection. PLoS Pathog 5:e1000524.

28. Bronnimann MP, Calton CM, Chiquette SF, Li S, Lu M, Chapman JA, Bratton KN, Schlegel AM, Campos SK. 2016. Furin Cleavage of L2 during Papillomavirus Infection: Minimal Dependence on Cyclophilins. J Virol 90:6224–6234.

29. Cerqueira C, Samperio Ventayol P, Vogeley C, Schelhaas M. 2015. Kallikrein-8 Proteolytically Processes Human Papillomaviruses in the Extracellular Space To Facilitate Entry into Host Cells. J Virol 89:7038–7052.

30. Richards KF, Bienkowska-Haba M, Dasgupta J, Chen XS, Sapp M. 2013. Multiple heparan sulfate binding site engagements are required for the infectious entry of human papillomavirus type 16. J Virol 87:11426–11437.

31. Richards RM, Lowy DR, Schiller JT, Day PM. 2006. Cleavage of the papillomavirus minor capsid protein, L2, at a furin consensus site is necessary for infection. Proc Natl Acad Sci U S A 103:1522–1527.

32. Surviladze Z, Dziduszko A, Ozbun MA. 2012. Essential roles for soluble virion-associated heparan sulfonated proteoglycans and growth factors in human papillomavirus infections. PLoS Pathog 8:e1002519.

33. DiGiuseppe S, Keiffer TR, Bienkowska-Haba M, Luszczek W, Guion LGM, Müller M, Sapp M. 2015. Topography of the Human Papillomavirus Minor Capsid Protein L2 during Vesicular Trafficking of Infectious Entry. J Virol 89:10442–10452.

34. Müller KH, Spoden GA, Scheffer KD, Brunnhöfer R, De Brabander JK, Maier ME, Florin L, Muller CP. 2014. Inhibition by cellular vacuolar ATPase impairs human papillomavirus uncoating and infection. Antimicrob Agents Chemother 58:2905–2911.

35. Xie J, Zhang P, Crite M, DiMaio D. 2020. Papillomaviruses Go Retro. Pathogens 9:267.

36. Bergant M, Banks L. 2013. SNX17 facilitates infection with diverse papillomavirus types. J Virol 87:1270–1273.

37. Day PM, Thompson CD, Schowalter RM, Lowy DR, Schiller JT. 2013. Identification of a role for the trans-Golgi network in human papillomavirus 16 pseudovirus infection. J Virol 87:3862–3870.

38. Lipovsky A, Popa A, Pimienta G, Wyler M, Bhan A, Kuruvilla L, Guie M-A, Poffenberger AC, Nelson CDS, Atwood WJ, DiMaio D. 2013. Genome-wide siRNA screen identifies the retromer as a cellular entry factor for human papillomavirus. Proc Natl Acad Sci U S A 110:7452–7457.

39. Popa A, Zhang W, Harrison MS, Goodner K, Kazakov T, Goodwin EC, Lipovsky A, Burd CG, DiMaio D. 2015. Direct binding of retromer to human papillomavirus type 16 minor capsid protein L2 mediates endosome exit during viral infection. PLoS Pathog 11:e1004699.

40. Speckhart K, Choi J, DiMaio D, Tsai B. 2024. The BICD2 dynein cargo adaptor binds to the HPV16 L2 capsid protein and promotes HPV infection. PLoS Pathog 20:e1012289.

41. Xie J, Heim EN, Crite M, DiMaio D. 2020. TBC1D5-Catalyzed Cycling of Rab7 Is Required for Retromer-Mediated Human Papillomavirus Trafficking during Virus Entry. Cell Rep 31:107750.

42. Campos SK. 2017. Subcellular Trafficking of the Papillomavirus Genome during Initial Infection: The Remarkable Abilities of Minor Capsid Protein L2. Viruses 9:370.

43. Lai K-Y, Rizzato M, Aydin I, Villalonga-Planells R, Drexler HCA, Schelhaas M. 2021. A Ran-binding protein facilitates nuclear import of human papillomavirus type 16. PLOS Pathog 17:e1009580.

44. Aydin I, Weber S, Snijder B, Samperio Ventayol P, Kühbacher A, Becker M, Day PM, Schiller JT, Kann M, Pelkmans L, Helenius A, Schelhaas M. 2014. Large scale RNAi reveals the requirement of nuclear envelope breakdown for nuclear import of human papillomaviruses. PLoS Pathog 10:e1004162.

45. Pyeon D, Pearce SM, Lank SM, Ahlquist P, Lambert PF. 2009. Establishment of human papillomavirus infection requires cell cycle progression. PLoS Pathog 5:e1000318.

46. Day PM, Weisberg AS, Thompson CD, Hughes MM, Pang YY, Lowy DR, Schiller JT. 2019. Human Papillomavirus 16 Capsids Mediate Nuclear Entry during Infection. J Virol 93:e00454–19.

47. Day PM, Thompson CD, Weisberg AS, Schiller JT. 2025. The COPII Transport Complex Participates in HPV16 Infection. Viruses 17:616.

48. DiGiuseppe S, Luszczek W, Keiffer TR, Bienkowska-Haba M, Guion LGM, Sapp MJ. 2016. Incoming human papillomavirus type 16 genome resides in a vesicular compartment throughout mitosis. Proc Natl Acad Sci U S A 113:6289–6294.

49. Day PM, Thompson CD, Schowalter RM, Lowy DR, Schiller JT. 2013. Identification of a Role for the *trans* -Golgi Network in Human Papillomavirus 16 Pseudovirus Infection. J Virol 87:3862–3870.

50. Woo T-T, Takeo Y, Harwood MC, Houck ET, DiMaio D, Tsai B. 2025. The nuclear import receptor importin-7 targets HPV from the Golgi to the nucleus to promote infection. Sci Adv 11:eadz6792.

51. Zhang W, Kazakov T, Popa A, DiMaio D. 2014. Vesicular Trafficking of Incoming Human Papillomavirus 16 to the Golgi Apparatus and Endoplasmic Reticulum Requires γ-Secretase Activity. mBio 5:e01777–14.

52. DiGiuseppe S, Luszczek W, Keiffer TR, Bienkowska-Haba M, Guion LGM, Sapp MJ. 2016. Incoming human papillomavirus type 16 genome resides in a vesicular compartment throughout mitosis. Proc Natl Acad Sci 113:6289–6294.

53. Mikuličić S, Strunk J, Florin L. 2021. HPV16 Entry into Epithelial Cells: Running a Gauntlet. Viruses 13:2460.

54. Nagashima S, Tábara L-C, Tilokani L, Paupe V, Anand H, Pogson JH, Zunino R, McBride HM, Prudent J. 2020. Golgi-derived PI ( 4 ) P-containing vesicles drive late steps of mitochondrial division. Science 367:1366–1371.

55. Kim Y, Burd CG. 2023. Lipid Sorting and Organelle Identity. Cold Spring Harb Perspect Biol 15:a041397.

56. Bankaitis VA, Garcia-Mata R, Mousley CJ. 2012. Golgi Membrane Dynamics and Lipid Metabolism. Curr Biol 22:R414–R424.

57. Keiffer TR, DiGiuseppe S, Guion L, Bienkowska-Haba M, Zwolinska K, Siddiqa A, Kushwaha A, Sapp MJ. 2025. HPV16 entry requires dynein for minus-end transport and utilizes kinesin Kif11 for plus-end transport along microtubules during mitosis. J Virol 99:e0093724.

58. Zádori Z, Szelei J, Lacoste MC, Li Y, Gariépy S, Raymond P, Allaire M, Nabi IR, Tijssen P. 2001. A viral phospholipase A2 is required for parvovirus infectivity. Dev Cell 1:291–302.

59. Ellis MV, James SR, Perisic O, Downes CP, Williams RL, Katan M. 1998. Catalytic Domain of Phosphoinositide-specific Phospholipase C (PLC). J Biol Chem 273:11650–11659.

60. Kanemaru K, Nakamura Y. 2023. Activation Mechanisms and Diverse Functions of Mammalian Phospholipase C. Biomolecules 13:915.

61. Bahk YY, Song H, Baek SH, Park BY, Kim H, Ryu SH, Suh P-G. 1998. Localization of two forms of phospholipase C-β1, a and b, in C6Bu-1 cells. Biochim Biophys Acta BBA - Lipids Lipid Metab 1389:76–80.

62. 2005. Nuclear phosphoinositide specific phospholipase C (PI-PLC)-ß1: a central intermediary in nuclear lipid-dependent signal transduction. Histol Histopathol 1251–1260.

63. Martelli AM, Follo MY, Evangelisti C, Falà F, Fiume R, Billi AM, Cocco L. 2005. Nuclear inositol lipid metabolism: More than just second messenger generation? J Cell Biochem 96:285–292.

64. Klein C, Gensburger C, Freyermuth S, Nair BC, Labourdette G, Malviya AN. 2004. A 120 kDa Nuclear Phospholipase Cγ1 Protein Fragment Is Stimulated in Vivo by EGF Signal Phosphorylating Nuclear Membrane EGFR. Biochemistry 43:15873–15883.

65. Yagisawa H, Okada M, Naito Y, Sasaki K, Yamaga M, Fujii M. 2006. Coordinated intracellular translocation of phosphoinositide-specific phospholipase C-δ with the cell cycle. Biochim Biophys Acta BBA - Mol Cell Biol Lipids 1761:522–534.

66. Okada M, Ishimoto T, Naito Y, Hirata H, Yagisawa H. 2005. Phospholipase Cδ_1_ associates with importin β1 and translocates into the nucleus in a Ca^2+^ -dependent manner. FEBS Lett 579:4949–4954.

67. Liu N, Fukami K, Yu H, Takenawa T. 1996. A New Phospholipase C δ4 Is Induced at S-phase of the Cell Cycle and Appears in the Nucleus. J Biol Chem 271:355–360.

68. Martelli AM, Billi AM, Manzoli L, Faenza I, Aluigi M, Falconi M, De Pol A, Gilmour RS, Cocco L. 2000. Insulin selectively stimulates nuclear phosphoinositide-specific phospholipase C (PI-PLC) β1 activity through a mitogen-activated protein (MAP) kinase-dependent serine phosphorylation. FEBS Lett 486:230–236.

69. Klein BM, Andrews JB, Bannan BA, Nazario-Toole AE, Jenkins TC, Christensen KD, Oprisan SA, Meyer-Bernstein EL. 2008. Phospholipase C beta 4 in mouse hepatocytes: Rhythmic expression and cellular distribution. Comp Hepatol 7:8.

70. Ackermann EJ, Conde-Frieboes K, Dennis EA. 1995. Inhibition of macrophage Ca(2+)-independent phospholipase A2 by bromoenol lactone and trifluoromethyl ketones. J Biol Chem 270:445–450.

71. Abayasekara RE, Flint AP. 1993. A novel phospholipase C inhibitor U73122, inhibits phospholipase C-independent processes in rat luteal cells. Biochem Soc Trans 21:353S.

72. Zhang XH, Zhao C, Seleznev K, Song K, Manfredi JJ, Ma ZA. 2006. Disruption of G1-phase phospholipid turnover by inhibition of Ca2+-independent phospholipase A2 induces a p53-dependent cell-cycle arrest in G1 phase. J Cell Sci 119:1005–1015.

73. Movahedi Naini S, Sheridan AM, Force T, Shah JV, Bonventre JV. 2015. Group IVA Cytosolic Phospholipase A_2_ Regulates the G_2_ -to-M Transition by Modulating the Activity of Tumor Suppressor SIRT2. Mol Cell Biol 35:3768–3784.

74. Hou C, Kirchner T, Singer M, Matheis M, Argentieri D, Cavender D. 2004. In vivo activity of a phospholipase C inhibitor, 1-(6-((17beta-3-methoxyestra-1,3,5(10)-trien-17-yl)amino)hexyl)-1H-pyrrole-2,5-dione (U73122), in acute and chronic inflammatory reactions. J Pharmacol Exp Ther 309:697–704.

75. Wang JW, Roden RBS. 2013. L2, the minor capsid protein of papillomavirus. Virology 445:175–186.

76. Lai K-Y, Rizzato M, Aydin I, Villalonga-Planells R, Drexler HCA, Schelhaas M. 2021. A Ran-binding protein facilitates nuclear import of human papillomavirus type 16. PLoS Pathog 17:e1009580.

77. Uhlorn BL, Jackson R, Li S, Bratton SM, Van Doorslaer K, Campos SK. 2020. Vesicular trafficking permits evasion of cGAS/STING surveillance during initial human papillomavirus infection. PLoS Pathog 16:e1009028.

78. Calton CM, Bronnimann MP, Manson AR, Li S, Chapman JA, Suarez-Berumen M, Williamson TR, Molugu SK, Bernal RA, Campos SK. 2017. Translocation of the papillomavirus L2/vDNA complex across the limiting membrane requires the onset of mitosis. PLoS Pathog 13:e1006200.

79. Hamdi M, Al-Matwi M, Elghoul N, Al-Kuwari H, Sayed TS, Riguene E, Nomikos M. 2025. Mammalian PI-Phospholipase C Isozymes: Structural and Functional Insights and Roles in Health and Disease. Medicina (Mex) 61:1054.

80. Suh P-G, Park J-I, Manzoli L, Cocco L, Peak JC, Katan M, Fukami K, Kataoka T, Yun S, Ryu SH. 2008. Multiple roles of phosphoinositide-specific phospholipase C isozymes. BMB Rep 41:415–434.

81. Bergeland T, Widerberg J, Bakke O, Nordeng TW. 2001. Mitotic partitioning of endosomes and lysosomes. Curr Biol 11:644–651.

82. Birky CW. 1983. The Partitioning of Cytoplasmic Organelles at Cell Division, p. 49–89. In Aspects of Cell Regulation. Elsevier.

83. Rabouille C, Jokitalo E. 2003. Golgi apparatus partitioning during cell division (Review). Mol Membr Biol 20:117–127.

84. Panagaki D, Croft JT, Keuenhof K, Larsson Berglund L, Andersson S, Kohler V, Büttner S, Tamás MJ, Nyström T, Neutze R, Höög JL. 2021. Nuclear envelope budding is a response to cellular stress. Proc Natl Acad Sci U S A 118:e2020997118.

85. Mettenleiter TC, Müller F, Granzow H, Klupp BG. 2013. The way out: what we know and do not know about herpesvirus nuclear egress: Herpesvirus nuclear egress. Cell Microbiol 15:170–178.

86. Bubeck A, Wagner M, Ruzsics Z, Lötzerich M, Iglesias M, Singh IR, Koszinowski UH. 2004. Comprehensive mutational analysis of a herpesvirus gene in the viral genome context reveals a region essential for virus replication. J Virol 78:8026–8035.

87. Fuchs W, Klupp BG, Granzow H, Osterrieder N, Mettenleiter TC. 2002. The interacting UL31 and UL34 gene products of pseudorabies virus are involved in egress from the host-cell nucleus and represent components of primary enveloped but not mature virions. J Virol 76:364–378.

88. Klupp BG, Granzow H, Mettenleiter TC. 2000. Primary envelopment of pseudorabies virus at the nuclear membrane requires the UL34 gene product. J Virol 74:10063–10073.

89. Reynolds AE, Ryckman BJ, Baines JD, Zhou Y, Liang L, Roller RJ. 2001. U(L)31 and U(L)34 proteins of herpes simplex virus type 1 form a complex that accumulates at the nuclear rim and is required for envelopment of nucleocapsids. J Virol 75:8803–8817.

90. Zeev-Ben-Mordehai T, Weberruß M, Lorenz M, Cheleski J, Hellberg T, Whittle C, El Omari K, Vasishtan D, Dent KC, Harlos K, Franzke K, Hagen C, Klupp BG, Antonin W, Mettenleiter TC, Grünewald K. 2015. Crystal Structure of the Herpesvirus Nuclear Egress Complex Provides Insights into Inner Nuclear Membrane Remodeling. Cell Rep 13:2645–2652.

91. Klupp BG, Granzow H, Fuchs W, Keil GM, Finke S, Mettenleiter TC. 2007. Vesicle formation from the nuclear membrane is induced by coexpression of two conserved herpesvirus proteins. Proc Natl Acad Sci U S A 104:7241–7246.

92. Roller RJ, Haugo AC, Kopping NJ. 2011. Intragenic and extragenic suppression of a mutation in herpes simplex virus 1 UL34 that affects both nuclear envelope targeting and membrane budding. J Virol 85:11615–11625.

93. Vu A, White S, Cassmann T, Roller RJ. 2021. Herpes Simplex Virus 1 UL34 Mutants That Affect Membrane Budding Regulation and Nuclear Lamina Disruption. J Virol 95:e0087321.

94. Florin L, Becker KA, Lambert C, Nowak T, Sapp C, Strand D, Streeck RE, Sapp M. 2006. Identification of a dynein interacting domain in the papillomavirus minor capsid protein l2. J Virol 80:6691–6696.

95. Pim D, Broniarczyk J, Siddiqa A, Massimi P, Banks L. 2021. Human Papillomavirus 16 L2 Recruits both Retromer and Retriever Complexes during Retrograde Trafficking of the Viral Genome to the Cell Nucleus. J Virol 95:e02068–20.

96. Cruz L, Biryukov J, Conway M, Meyers C. 2015. Cleavage of the HPV16 Minor Capsid Protein L2 during Virion Morphogenesis Ablates the Requirement for Cellular Furin during De Novo Infection. Viruses 7:5813–5830.

97. Kieback E, Müller M. 2006. Factors influencing subcellular localization of the human papillomavirus L2 minor structural protein. Virology 345:199–208.

98. Wang JW, Roden RBS. 2013. L2, the minor capsid protein of papillomavirus. Virology 445:175–186.

99. Peng W-Y, Abere B, Shi H, Toland S, Smithgall TE, Moore PS, Chang Y. 2023. Membrane-bound Merkel cell polyomavirus middle T protein constitutively activates PLCγ1 signaling through Src-family kinases. Proc Natl Acad Sci U S A 120:e2316467120.

100. Broniarczyk J, Massimi P, Pim D, Bergant Marušič M, Myers MP, Garcea RL, Banks L. 2019. Phosphorylation of Human Papillomavirus Type 16 L2 Contributes to Efficient Virus Infectious Entry. J Virol 93:e00128–19.

101. Bronnimann MP, Chapman JA, Park CK, Campos SK. 2013. A Transmembrane Domain and GxxxG Motifs within L2 Are Essential for Papillomavirus Infection. J Virol 87:464–473.

102. Oh C, Buckley PM, Choi J, Hierro A, DiMaio D. 2023. Sequence-independent activity of a predicted long disordered segment of the human papillomavirus type 16 L2 capsid protein during virus entry. Proc Natl Acad Sci 120:e2307721120.

103. Bienkowska-Haba M, Luszczek W, Keiffer TR, Guion LGM, DiGiuseppe S, Scott RS, Sapp M. 2017. Incoming human papillomavirus 16 genome is lost in PML protein-deficient HaCaT keratinocytes. Cell Microbiol 19.

104. Guion L, Bienkowska-Haba M, DiGiuseppe S, Florin L, Sapp M. 2019. PML nuclear body-residing proteins sequentially associate with HPV genome after infectious nuclear delivery. PLoS Pathog 15:e1007590.

105. Dorosz K, Majewska L, Kijowski J. 2025. Structure and Function of PML Nuclear Bodies: A Brief Overview of Key Cellular Roles. Biomolecules 15:1291.

106. Uggè M, Simoni M, Fracassi C, Bernardi R. 2022. PML isoforms: a molecular basis for PML pleiotropic functions. Trends Biochem Sci 47:609–619.

107. Ferguson BJ, Dovey CL, Lilley K, Wyllie AH, Rich T. 2007. Nuclear phospholipase C gamma: punctate distribution and association with the promyelocytic leukemia protein. J Proteome Res 6:2027–2032.

108. Schweiger L, Lelieveld-Fast LA, Mikuličić S, Strunk J, Freitag K, Tenzer S, Clement AM, Florin L. 2022. HPV16 Induces Formation of Virus-p62-PML Hybrid Bodies to Enable Infection. Viruses 14:1478.

109. Maa MC, Leu TH, Trandel BJ, Chang JH, Parsons SJ. 1994. A protein that is highly related to GTPase-activating protein-associated p62 complexes with phospholipase C gamma. Mol Cell Biol 14:5466–5473.

110. Buck CB, Pastrana DV, Lowy DR, Schiller JT. 2004. Efficient intracellular assembly of papillomaviral vectors. J Virol 78:751–757.

111. Buck CB, Pastrana DV, Lowy DR, Schiller JT. 2005. Generation of HPV pseudovirions using transfection and their use in neutralization assays. Methods Mol Med 119:445–462.

112. Buck CB, Thompson CD, Pang Y-YS, Lowy DR, Schiller JT. 2005. Maturation of papillomavirus capsids. J Virol 79:2839–2846.

113. Buck CB, Thompson CD. 2007. Production of papillomavirus-based gene transfer vectors. Curr Protoc Cell Biol Chapter 26:Unit 26.1.

114. Brown LY, Dong W, Kantor B. 2020. An Improved Protocol for the Production of Lentiviral Vectors. STAR Protoc 1:100152.

115. DiGiuseppe S, Luszczek W, Keiffer TR, Bienkowska-Haba M, Guion LGM, Sapp MJ. 2016. Incoming human papillomavirus type 16 genome resides in a vesicular compartment throughout mitosis. Proc Natl Acad Sci U S A 113:6289–6294.

